# Chronic stress abnormalize microglial-vascular interaction via AT1R signaling and microglia activation to drive depression-like behaviors

**DOI:** 10.64898/2025.12.24.696453

**Authors:** Hang Gao, Zeyuan Ding, Biao Xu, Yuqing Wu, Meizhen Zhu, Xiaoyue Zhang, Fujian Qi, Junru Liu, Quan Fang, Yanli Ran

## Abstract

Depression entails a range of pathological processes in the brain, typically characterized by microglia activation, neural dysfunction, and vascular dysregulation. While it is well established that activated microglia inflict damage to neurons through direct and indirect microglia-neuron interactions, their interaction with vasculature and the associated regulation of vascular structure and function in depression remain poorly understood. Here, using a mouse model of depression induced by chronic unpredictable stress (CUS), we find that in stressed mice, activated microglia present reduced motility and decreased contact area with vasculature, which is accompanied by pathological vascular structure and function, marked by reduced vessel density, narrowed vascular diameters, and sluggish blood flow velocity. We observe that vascular structure in the stressed mice is rapidly modified by the vascular-contacted microglia. These vascular-contacted microglia show upregulated angiotensin II type 1 receptor (AT1R) expression, which is not present in the vascular-uncontacted microglia. Furthermore, we demonstrate that AT1R signaling together with microglia activation under chronic stress are critical mediators for the altered microglial-vascular contact situation, vascular dysregulation, as well as the depression-like behaviors. Together, our findings reveal that under CUS, the aberrant microglial-vascular interaction, including the dysregulated microglial vasoregulation arising from AT1R signaling and microglia activation, leads to depression, forging a new link between chronic stress and depression.

## 1 Introduction

Chronic stress is a main risk factor for psychiatric diseases like depression. It drives a range of pathological processes in the brain, including neuroinflammation, neural impairments and cerebral hypoperfusion ^1–4^. These pathological processes are monitored and modified by the immune cell in the central nervous system, the microglia ^5–8^. Sustained exposure to stress triggers the activation of microglia ^7,9,10^. The activated microglia promote the production and release of pro-inflammatory cytokines and other neurotoxic factors, further exacerbating the brain disorders ^11–13^. Thus, chronic stress induces depression is commonly thought to be resulted from the inflammation-related mechanisms ^14–16^. However, in addition to mediating inflammation, microglia also engage in direct interactions with surrounding cells and perform local regulatory functions, such as modulating synaptic pruning through microglia-neuron interactions ^17,18^. In the brain, microglial processes are recruited to blood vessels and the loss of microglial-vascular contact results in abnormal vascular diameter and blood perfusion ^19–22^, suggesting close interactions between microglia and vasculature. In many pathologies, including depression, vascular dysregulation is known to occur. Given the essential roles of the vascular system in the brain, understanding microglial-vascular interactions and the associated effects on vascular properties is vital for our comprehensive understanding of the mechanisms underlying vascular dysregulation and the pathogenesis of depression.

Previous studies have reported that microglia can influence vascular physiology via the release of potent signalling molecules and the expression of vasoactive genes ^20,21,23–25^. Reciprocally, the vascular components and phenotype shape microglial behaviors. For instance, endothelial cells release chemokines (e.g., CX3CL1) that influence microglial activation states and inflammatory responses via receptors such as CX3CR1 ^26–28^. When the vasculature changes, microglia exhibit rapid responses. It has been reported that microglia migrate to the leaked blood-brain barrier (BBB) position and contribute to the repairment of BBB integrity ^29^. Additionally, following transient ischemia, the processes’ motility of microglia in contact with vasculature correlates with the reduction of blood flow velocity ^30^. In depression, both the activation of microglia and the deficit of cerebral blood perfusion occur ^1,14^, suggesting altered interactions between microglia and vasculature. However, a systematic survey of microglial-vascular interaction in the context of depression and the underlying mechanisms is still lacking.

In the current study, we exploit the microglial-vascular interaction in the context of CUS-induced depression and the associated effects on depression-like behaviors. We found that in CUS-induced depression, microglia altered their morphodynamics and reduced their contact area with the vasculature. Concurrently, after CUS, the vasculature exhibited structural and functional compromises, which was rapidly regulated by the vascular contacted microglia. Furthermore, we demonstrated that the elevated AT1R signaling in microglia and the activation of microglia under CUS served as key mediators for the aberrant microglial-vascular interaction that resulted in dysregulation of vascular structure and function and finally depression-like behaviors, providing a novel mechanistic basis for the contribution of microglial-vascular interaction to chronic stress-induced depression, which might guide new therapies targeting depression and other vascular pathologies.

## 2 Materials and methods

### Animals

The C57BL/6 mice used in this study were procured from the Experimental Animal Centre of Lanzhou University. The CX3CR1^GFP/GFP^ mice were originally purchased from the Jackson Laboratory by Prof. Shengxiang Zhang, who generously gave them to us. The heterozygous CX3CR1^GFP/+^ mice were generated through the crossbreeding of homozygous CX3CR1^GFP/GFP^ and C57BL/6 mice. Mice aged 8-12 weeks, with a body weight ranging from 18 to 22 grams were used throughout this study. All animals were maintained in the animal facility of Lanzhou University under controlled conditions (temperature, humidity, and 12-h light/dark cycle), with free access to food and water. The entire experimental protocol was performed in strict compliance with the European Community Guidelines (2010/63/EU) and ARRIVE Guidelines, and was approved by the Ethics Committee of Lanzhou University.

### Chronic unpredictable stress procedure

The experimental mice were randomly allocated to either the chronic unpredictable stress (CUS) group or the control group. The mice in the control group were maintained under normal housing conditions, whereas those in the CUS group were subjected to three types of stress treatments daily for a period of 10 days and were housed in individual cages throughout the treatment cycle. Three stressors were randomly selected each day from the following proven and approved standard stressors to ensure unpredictability for the subjects: 4-hour tilt cage without bedding (at an angle of 45°), 24-hour wet bedding, 24-hour constant illumination, 2-hour crowding (10 mice per cage), 4-hour social isolation (Use 15*15 cm small boxes to isolate the mice), 30-minute elevated platform exposure, 2-hour restraint behavior, and 2-hour rapid light on-off (with an interval of 7 minutes). Water and food were freely available throughout the entire CUS procedure. The control mice were kept under standard conditions for ten days with free access to food and water.

### Drug treatment

***Candesartan treatment.*** A stock solution of 40 mg/ml was prepared by dissolving 2 mg of candesartan (TargetMol, CV-11974, USA) in 50 μl of DMSO (Sigma-Aldrich, 276855, USA). Subsequently, 50 μl of the stock solution was mixed with 300 μl of PEG300 (Solarbio, IP9020, China), followed by the addition of 50 μl of Tween 80 (Solarbio, IP9020, China), and finally, 600 μl of ddH₂O (Servicebio, G4700, China) were added to obtain a working solution with a concentration of 2 mg/ml. Mice were administered the drug or vehicle via intraperitoneal (i.p.) injections at a dosage of 2 mg/kg every day starting from the first day of CUS.

***Minocycline treatment.*** After 5 days of the CUS procedure, minocycline (Mino, 50 mg/kg, 2 mg/ml, Macklin, M812977, China) or physiological saline (Sigma-Aldrich, 52455, USA) was injected intraperitoneally. The injections were administered once per day between 11:00 and 11:30 in the morning.

### Behavioral testing

***Forced swimming test (FST).*** Mice were placed in a transparent glass cylinder (height: 25 cm, diameter: 10 cm) filled with water (depth: 10 cm, temperature: 25 ± 1°C) for 6 minutes. Immobility was defined as the mouse floating passively without struggling, exhibiting only necessary movements to maintain head above the water surface. Mouse behaviors were video-recorded, and the immobility duration during the last 4 minutes of the test was analyzed.

***Tail suspension test (TST).*** Mice were suspended 50 cm above the floor via adhesive tape affixed 1 cm from the tail tip for 6 minutes. Immobility was defined as passive hanging with complete motionlessness. Mice that climbed their tails were excluded from subsequent analysis. The immobility duration during the last 4 minutes of the suspension period was analyzed.

***Electronic von Frey.*** The mechanical threshold of the mice was assessed using an electro-mechanical prickling apparatus. Mice were placed in a transparent plastic box (8 × 8 × 5.5 cm) with a metal mesh at the bottom 2 days before the start of the normal experiment and acclimated for at least 30 min a day to minimize the influence of the stress response on the experimental results. Before the experiment, the electronic mechanical stabbing instrument was connected to the data acquisition system and calibrated to ensure that the measurement accuracy reached 0.2 g. For mechanical paw withdrawal threshold (PWT) measurement, von Frey filaments were applied vertically to the right hind paw of the mouse, and the stimulus intensity was slowly increased until the mouse exhibited a significant paw withdrawal response. The force value at this time was recorded, which was the paw withdrawal threshold.

### Immunofluorescence

The mice were deeply anaesthetized with ethyl carbamate solution (at a dosage of 1 g/kg body weight) via intraperitoneal injection and then intracardially perfused with 0.01 M phosphate buffered saline (PBS, BBI, B640011, China) and 4% paraformaldehyde (PFA, BBI, E672002, China). The brain tissue was removed and fixed overnight at 4°C with 4% PFA. Subsequently, the brain sections were sliced to a thickness of 30 μm using a Leica vibrating-blade microtome in 0.01 M PBS and stored in PBS solution. Three to four brain slices were selected from each mouse and placed in 24-well plates, followed by three washes with PBS for 15 minutes each. Subsequently, sections were blocked for 2 h in PBS (5% goat serum, 0.3% Triton X-100). Post-blocking, sections were incubated with rabbit anti-Iba1 (1:1000, Wako, AB_839504) overnight at 4°C. After primary antibody incubation, sections were washed with PBS (3×15 min), then incubated with goat anti-rabbit secondary antibody (1:1000, Alexa Fluor 488, Invitrogen, AB_143165) for 2 h at room temperature in dark. Finally, after incubation, the slides were washed three additional times with PBS for 15 minutes each, and all the sections were mounted with antifade mounting medium (Beyotime, P0126, China). Images were collected using a Nikon laser scanning confocal microscope (Nikon AX R MP), and the cell density and morphology of cerebral cortex microglia under different conditions were analyzed using ImageJ software.

### RNAscope in situ hybridization

Mouse brain tissues were sectioned into 15 μm slices using a freezing microtome (Leica, CM1860, Germany). Endogenous peroxidase was quenched with RNAscope hydrogen peroxide solution (ACD) for 10 min, followed by incubation with RNAscope protease III (ACD) at 40 °C for 30 min. Sections were then incubated with probe mixture at 40 °C for 2 h. C1 probes included Pecam1 (ACD, Cat: 316721) and Angpt2 (ACD, Cat: 406091), labelled with Opal 570 fluorophore (1:750) to detect the mRNA expression of CD31 and angiotensin II (AngII). C2 probe Agtr1a (angiotensin II receptor – AT1R, ACD, Cat: 4881161-C2) was labelled with Opal 690 fluorophore (1:750). After probe hybridization, sections were developed and subjected to routine immunofluorescence staining for microglia labeling.

### Vascular labelling using gelatin-based lipophilic dye solution (VALID)

The blood vessels in fixed brain tissue were labelled using VALID as previously described ^31^. Initially, 60 mg of DiI powder (Aladdin, D131225, China) was dissolved in 10 ml of anhydrous ethanol to prepare a stock solution at a concentration of 6 mg/ml. This solution was stored at room temperature in the absence of light. For vascular labelling, Type A porcine skin gelatin (Sigma-Aldrich, V900863, USA) was dissolved in distilled water to prepare a 4% (w/v) gelatin solution. Subsequently, the DiI stock solution was mixed with the porcine skin gelatin solution at a ratio of 1:50 to prepare the working solution for vascular labelling. The prepared working solution was maintained in a 40℃ constant-temperature water bath until use. Experimental mice were anaesthetized via intraperitoneal injection of 1 g/kg ethyl carbamate. Following anaesthesia, 0.01 M PBS was perfused through the heart to remove residual blood, followed by perfusion of 10-15 ml of the prepared porcine skin gelatin working solution. After perfusion, the mouse body was stored overnight in a 4℃ refrigerator to facilitate gel solidification. Subsequently, the brain tissue was dissected and fixed in 4% paraformaldehyde for 24 hours. Following fixation, the brain tissue was sectioned into 100 µm thick slices using a Leica vibratome (Leica, VT1000S) and mounted with an anti-fluorescence quenching medium for preservation.

### Confocal imaging

Imaging of the sections was conducted using a Nikon laser scanning confocal microscope (Nikon AX R MP). For excitation, 488 nm light was used to stimulate the EGFP fluorescence in microglia cells, while 561 nm light was employed to excite the red fluorescence of diI-labelled blood vessels. A 25× water immersion objective with a numerical aperture (NA) of 1.10 was utilized, and 3D images were acquired at 1-micron intervals in the Z direction. The obtained images were processed using ImageJ for quantification of microglia density and process area. Three-dimensional reconstructions of microglia and blood vessels were performed using Imaris and ImageJ, enabling detailed morphological analysis. Additionally, 3D co-localization analysis between microglia and blood vessels was performed using Imaris.

### Two-photon imaging *in vivo*

***Preparation of two-photon imaging window.*** Prior to experimentation, all surfaces and equipment were disinfected using a 75% alcohol solution. After deep anaesthesia with isoflurane, the mice were fixed on the brain stereotaxic apparatus, and the heads were cleaned to expose the skull. A skull drill was used to draw a circular outline with a diameter of slightly more than 3 mm at AP1.0 mm and ML1.5 mm from the bregma point, and then the polished circular skull fragment was carefully separated from the skull with forceps. Next, the window was cleaned with antibiotics immediately, and the craniotomy site was kept moist throughout the procedure. When the cleaning was completed, a circular glass sheet with a diameter of 3 mm was placed on the window surface and fixed with tissue adhesive (3M Company, 1469SB, USA). Following the surgery, the mice were transferred to a warm recovery cage until they regained consciousness. For fourteen days post-surgery, 300 μl of antibiotics were administered intraperitoneally on daily.

***Two-photon imaging.*** *In vivo* imaging was performed four weeks after surgery, when the mice had fully recovered. The procedure utilized a Nikon two-photon laser scanning confocal microscope equipped with a 25× 1.10NA water immersion objective lens. The mice were securely fastened onto the awake mouse fixation mount, and during the entire imaging session, they remained under isoflurane anaesthesia to ensure stability and compliance. Dextran (Invitrogen, 2600119, USA) was intravenously administered into mice for the purpose of labelling blood vessels. Subsequently, recordings were captured using two-photon microscopy technology. The pulsed laser operated at a wavelength of 1000. The green and red channels were selectively engaged to image microglia and blood vessels respectively, with a precisely set z-step value of 1 µm. The imaging depth spanned from 90 to 160 µm, and imaging at two different magnifications, namely Zoom1 and Zoom5, was sequentially executed. Moreover, a single blood vessel was swiftly scanned 520 times to accurately measure the blood flow velocity. Meanwhile, individual microglia were imaged at 0 hours and 1 hour respectively to meticulously document their dynamic alterations. Upon the completion of imaging, ImageJ and Imaris software were employed to conduct comprehensive three-dimensional co-localization analyses of microglia and blood vessels, in-depth morphological investigations of microglia, detailed examinations of microglia’s dynamic changes, as well as precise measurements of blood vessel diameters. Additionally, the dedicated analysis software NIS-Elements incorporated within the Nikon two-photon microscope system was utilized to perform the blood flow velocity analysis, ensuring high precision and reliability throughout the entire process.

### Data processing

Imaris (Version 9.0.1, Bitplane AG) and ImageJ (Version 1.51n) were used for image analyzation and visualization.

***Cell counting.*** Perform manual cell counting using ImageJ (version 1.51n) and import the image into ImageJ, the "plugins-Analysis-Cell counter" module of ImageJ was used to manually count microglia.

***Microglial morphology.*** Microglial skeletonization was performed using ImageJ plugins. Briefly, images were processed with the "Plugin-Process-Smooth 3D" plugin (radius: 0.3-0.8). The total number of microglial process endpoints and total process length per cell were quantified using an analytical skeleton plugin (http://imagejdocu.tudor.lu), and skeletonized images were generated via the 2D/3D Skeletonization plugin.

***Vascular blood flow.*** In the experimental setup, the blood vessels were labeled with the fluorescent dye dextran (Invitrogen, 2600119, USA) via tail vein injection. Concurrently, the plasma was tagged with dextran, which caused it to appear red. Subsequently, the blood flow velocity was measured by employing the two-photon line-scanning module. In this process, the same cross-section of the blood vessel was repeatedly scanned with a laser. The individual images obtained from these scans were then integrated to form a composite image. For the purpose of velocity calculation, the X direction was defined as the displacement of red blood cells. In the Y direction, the time interval for each row corresponded to the time required for the line scan to complete one full repetition. Based on these definitions, the blood flow velocity could be computed using the following formula: Calculated speed = X distance / Y distance.

***Vascular density.*** The vessels were rendered on the surface of Imaris and then their 3D structures were reconstructed. During the reconstruction process, the threshold was adjusted to distinguish the vessels from the background. The volume of blood vessels was calculated after 3D reconstruction.

***Vascular diameter.*** 3D images of vascular segments were superposed to 2D pictures in the z-axis by maximum intensity projection in ImageJ. Images were calibrated by entering a scale directly related to the microscope settings used during image acquisition, and vessels were then analyzed for diameter and coverage area with the "Analytic-Measure" and "surface" modules.

### Colocalization analysis

***Microglial-vascular contacts.*** The two-photon acquired images were imported into Imaris and analyzed using the "colocalization" module. The blood vessels and microglia were distinguished into different color channels based on their distinct labelling. Then, a threshold was set to select the microglia and blood vessels. Finally, the volume and proportion of the co-localized regions were calculated.

***Morpho-dynamic changes of microglia.*** The area where the entire microglial processes were located was obtained using ImageJ software. First, a 30 μm stack of the same microglia from different imaging timepoints was selected and stacked in the z-axis. Then, images acquired at 0 h and 1 h were superimposed, after which the 0-h images were set to red pseudo-color and the 1-h images were set to green pseudo-color. On superimposed images at each of the two timepoints, yellow pixels indicate no movement of microglia, green pixels indicate microglial process extension, and red pixels indicate process retraction. After applying a unified threshold to the MIP images, the area occupied by the protrusions was obtained using the “Analyze particles” tool. The area of the extended and contracted pixels divided by the area of the stable pixels was used to determine the Motility index (Motility index = (A_retraction + A_extension)/A_stable).

***RNAscope data.*** Colocalization analysis of microglia, AngII, and AT1R was performed using ImageJ. First, the channels were split through the "Image-Color-split channels" module. Then adjust the threshold by the "Image-Adjust-Threshold" module, and then the "AND" operator is used to obtain two or multi-channel co-localization images. The "Creat mask" was used to form the co-labelled signal, and the "Analytic-Analyze particals" module was used to quantify the co-localization.

### Statistics

Statistical analyses were performed using GraphPad Prism 8.0.2. Two-tailed unpaired t-tests were used for two-group comparisons; one-way ANOVA followed by Tukey’s post hoc test was applied for multiple comparisons among three or more groups. Results were expressed as mean ± standard error of the mean (SEM), with statistical significance set at p < 0.05.

## 3 Results

### 3.1 Chronic stress induced depression associated with microglial morphodynamic changes

To induce depression in mice, we expose mice to CUS, which is a well-established protocol that induces depression-like behaviors ^32^. After CUS, we performed FST and TST in mice that underwent CUS to evaluate the behaviors of mice (Fig.1a-c). We found that male mice after CUS displayed depression-like behaviours, with increased immobility time in FST and TST (Fig.1b, c). Whereas, for female mice, the immobility time in FST and TST was not significantly affected by CUS experiences (sFig.1). Therefore, we used male mice throughout this study.

**Figure 1.**
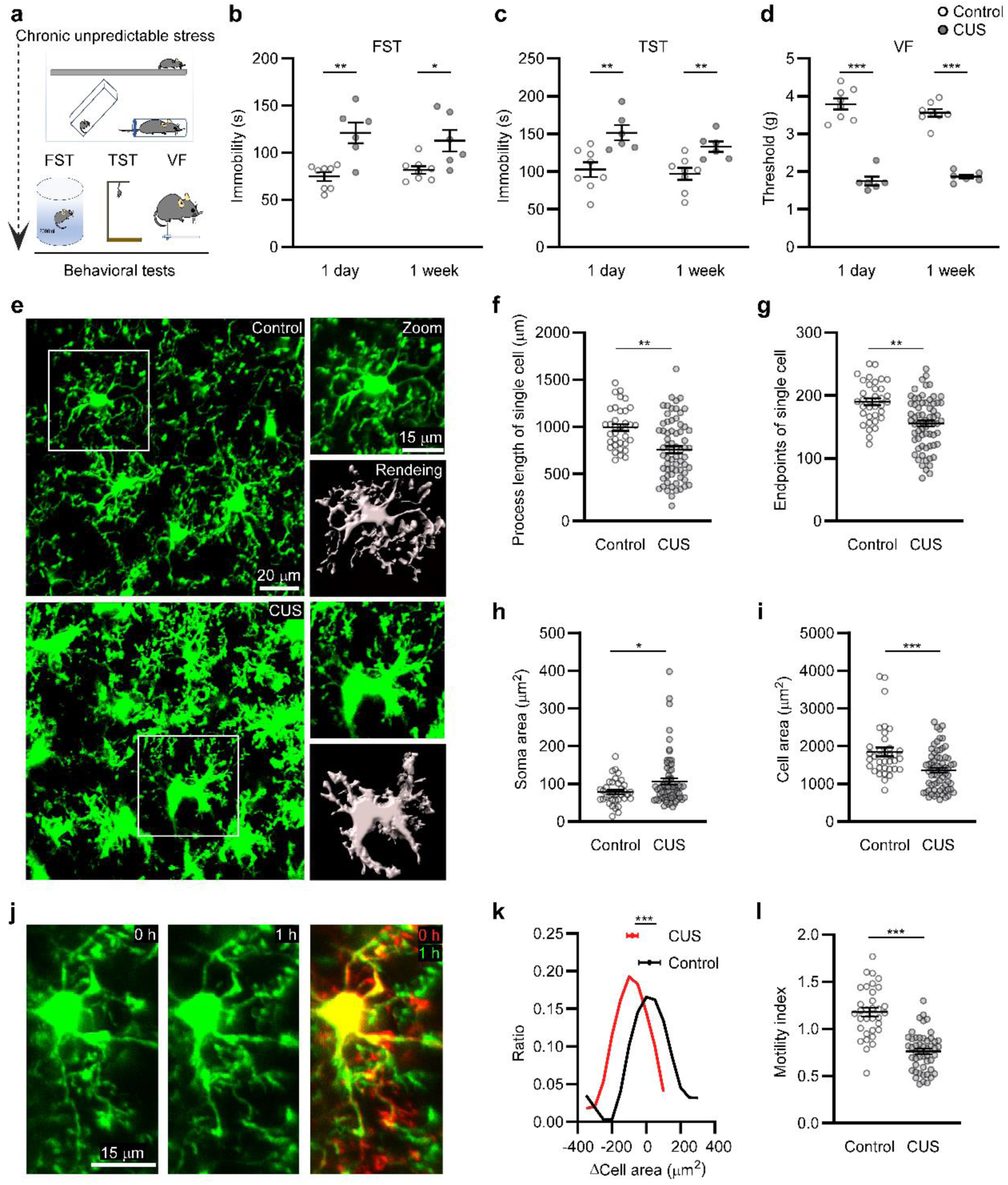
CUS leads to depressive-like behaviors and microglial morphodynamic changes. **(a)** Illustrating mice with CUS treatment and behavioral tests. **(b-d)** The immobility time of mice in FST, TST and paw withdrawal threshold of mice in von Frey test. Control: n = 8; CUS: n = 6 mice. **(e)** Representation of microglial morphology acquired from two-photon imaging *in vivo*. Zoomed images showing single microglia cropped from the white rectangle indicated in the left image (right up) and the corresponding rendered morphology using IMARIS (right down). **(f-i)** The analysis for total length, endpoints, soma size and cell area in single microglia. Control: n = 4/35, mice/slice; CUS: n = 6/67, mice/ cells. **(j)** Top-view of the same microglia at 0 h and 1 h after 10 days of CUS. **(k)** Histogram (curve) and averages (inset) of microglial area changes based on images in (j) (control: n = 4/33; CUS: n = 6/49, mice/cells). **(l)** Motility index of the same microglia analyzed in (k).

Besides the depression-like behaviors, mice presented hypersensitivity to mechanical stimuli (Fig.1d), suggesting sensory processing disorders accompanied by depression. Here, the primary somatosensory cortex – S1, an essential region for sensory processing, is very likely involved in this disturbed function ^33,34^. As microglia activation has been commonly regarded as a critical mediator for depression and the related functional disorders in the brain ^14,15^, we examined the microglial numbers and morphologies in S1, which were thought to relate to their activation status. We first checked these parameters in the fixed brain slices, we found that both the immuno-stained Iba1-positive cells and CX3CR1-GFP cells in CUS mice were significantly more than that from mice under normal conditions, and presented shorter process lengths and fewer endpoints in individual cells (sFig.2). We then semi-automatically measured more morphological parameters of individual microglia based on their 3D images acquired from CX3CR1^GFP/+^ mice *in vivo*. The *in vivo* data confirmed that microglia in mice that underwent CUS exhibited activated morphology, showing increased soma sizes, decreased process lengths, branch points and cell areas (Fig.1e-i). Furthermore, we found that the microglial morphology changed continuously after CUS. Even within one hour, microglia in CUS mice further decreased cell area, which was not observed in normal conditioned mice (Fig.1j, k). Additionally, the motile ability of microglia in CUS mice was significantly weaker than that of the normal mice (Fig.1l). Since microglia survey the surroundings using their motile processes ^5,6,8,10^, their dynamic morphology changes under CUS might be associated with altered detection ability and functional relationships with the surroundings.

**Figure 2.**
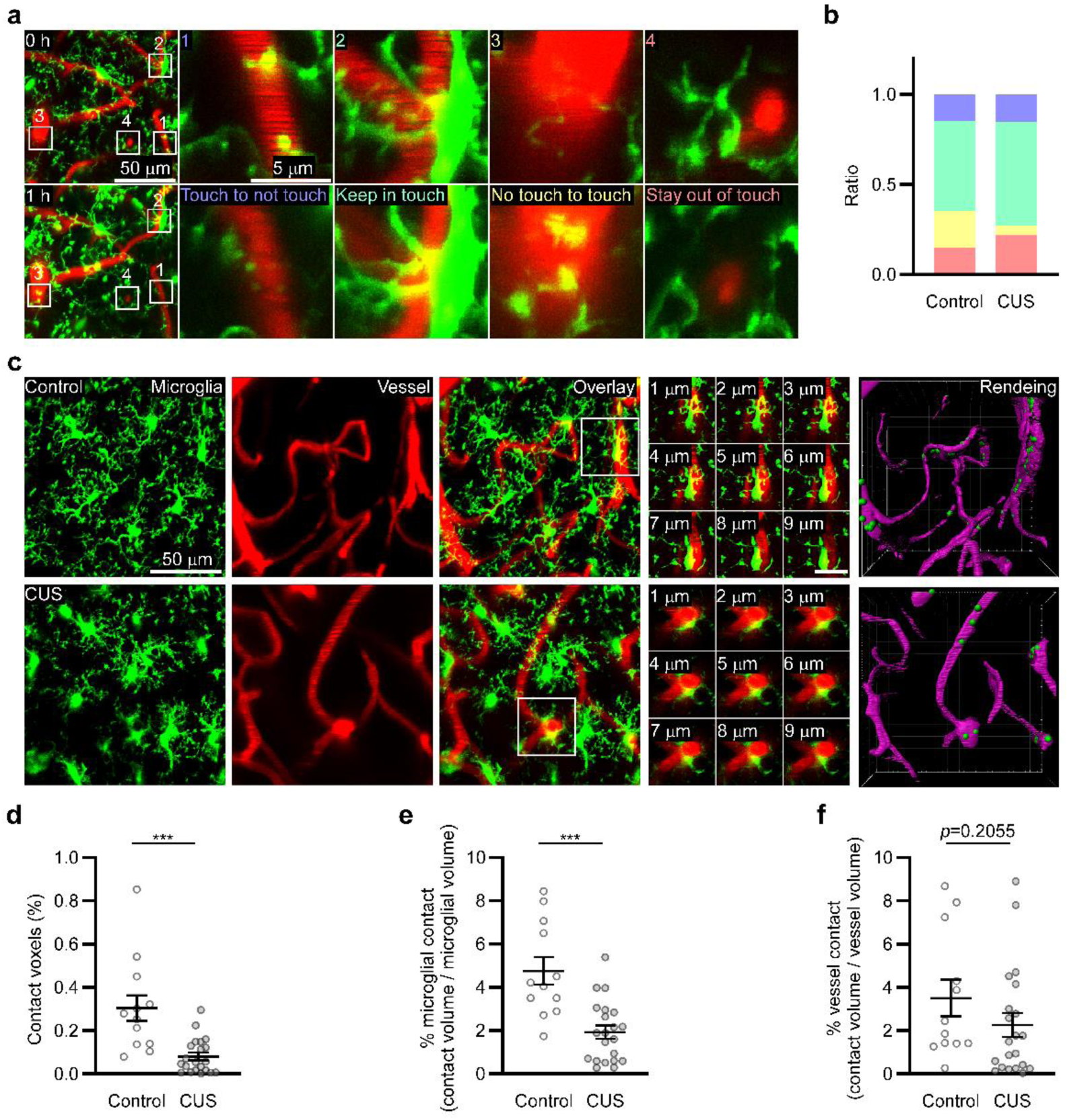
CUS induces a reduction in physical contacts between microglia and vasculature. **(a)** Z-projection of image stacks showing microglia (green) and blood vessels (red) in the same field acquired at two timepoints (0 h & 1 h) after CUS. Zoomed images showing different dynamic changes regarding microglial-vascular contacts at the two imaged timepoints, as indicated by the white rectangles in the left images. **(b)** Proportion of different microglia vascular contact situations over one hour (Control: 54 vascular segments in total, contact to no contact 0.15, maintain contact 0.50, no contact to contact 0.20, maintain no contact 0.15; CUS: 78 vascular segments in total, contact to no contact 0.15, maintain contact 0.58, no contact to contact 0.05, maintain no contact 0.22). **(c)** Exemplary 60-µm-thick *in vivo* two-photon images showing microglia (green) and the vasculature (red) in S1 with a rectangle in the overlay image indicating microglia vascular contacts. **(d)** Representative images showing microglia contacting the vasculature in a 9-µm-thick tissue volume, as indicated by the rectangle in (c). **(e)** Reconstructions of microglia (green) in contact with vasculature (magenta) generated from 3D two-photon images using IMARIS. **(f)** Ratio of microglial-vascular contact area to total volumetric area within the 3D-imaged region. **(g)** Percentile of microglial vascular contact area relative to total microglial area. **(h)** Percentile of microglial-vascular contact area relative to total vascular area within the 3D-imaged volume. d-e: Control: n = 4/12; CUS: n = 6/21 mice/slices.

### 3.2 Chronic stress leads to reduced contact between microglia and blood vessels

As microglia are known to contact with vasculature ^19–22^, we next examined whether the physical contact between microglia and vasculature changed following CUS. For this, we performed intracardial perfusion in deeply anaesthetised CX3CR1^GFP/+^ mice with a red dye as described recently ^31^. This allowed us to image both blood vessels and microglia under a microscope in the fixed brain slices. Images showed that microglia in contact with vasculature in the CUS mice were significantly less than those in the normal conditions (sFig.3a-c).

We further examined the microglial-vascular contacts in living mice. To fluorescently label blood vessels and microglia, we injected dextran via the tail vein in CX3CR1^GFP/+^ mice. Data acquired through *in vivo* two-photon imaging showed that, for both normal and CUS conditioned mice, there were a fraction of microglia in contact with vasculature with their soma and/or processes (Fig.2a). When quantitatively assessing these contacts, we found that microglia in CUS mice tend to lose their contact with blood vessels. Within one-hour, the possibility of microglia lost their contacts with vessels was three-fold of that newly established contacts with vessels (Fig.2b). However, in unstressed mice, the possibility of microglia that lost their contacts with vessels was similar to those newly formed contacts (Fig.2b). In line with these results, the contacts were strikingly less in chronically stressed mice than in normal conditioned mice, displaying a remarkably decreased contact areas and lower percentage of contacted microglia (Fig.2c-e). Notably, unlike the decreased total microglial-vascular contact areas, there were no obvious changes in the frequency of vessels receiving microglia contacts after CUS (Fig.2f), suggesting a reduction of vessel density in chronically stressed mice.

### 3.3 Abnormal vascular structure and function in chronically stressed mice

To investigate the detailed vascular alterations induced by CUS, we scanned the same area in S1 before and after CUS using two-photon imaging and longitudinally compared the vascular characteristics. As indicated above, after CUS, there was significantly lower vessel density compared with that before CUS (Fig.3a, b). Images from brain slices also showed that vascular density in chronically stressed mice was significantly lower than that in the normal conditioned mice(sFig.3d). The vascular density analyzations indicate that chronic stress induces deficits in cerebral blood perfusion, which is consistent with previous findings in depression ^1,35^.

**Figure 3.**
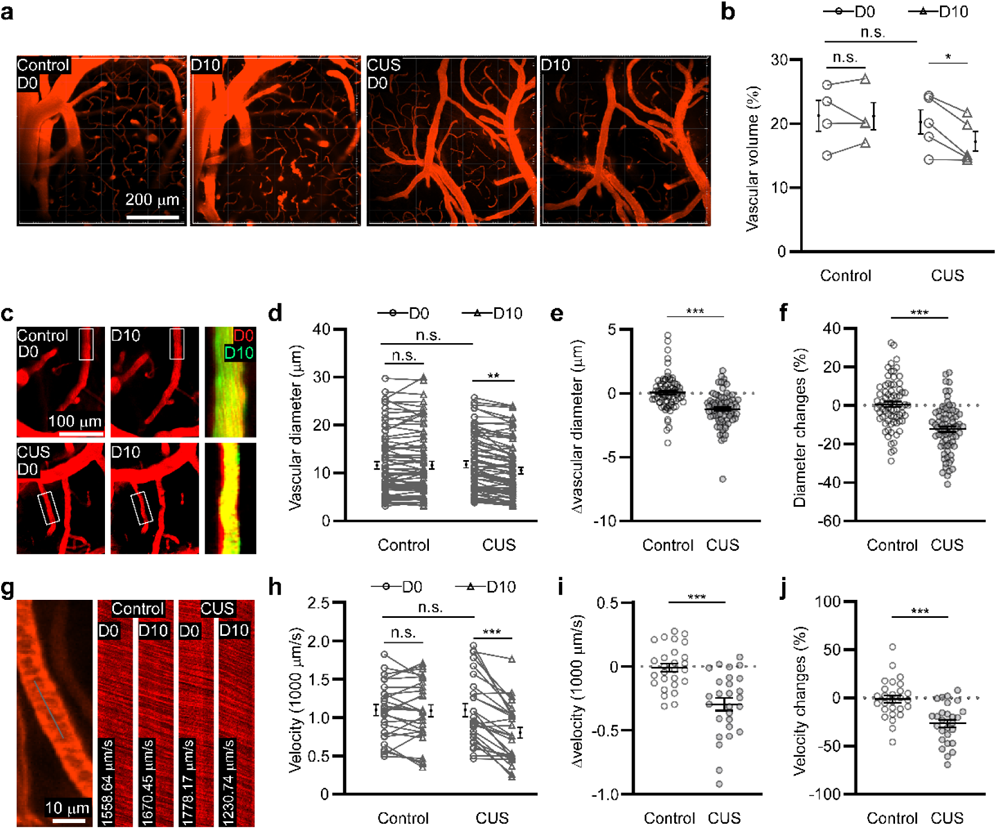
Vasculature exhibits pathological structural and functional properties following CUS. **(a)** Exemplary images showing vasculature from the same regions in S1 before (D0) and after (D10) CUS. **(b)** Quantification of vessel density (control: n = 4 mice; CUS: n = 5 mice). **(c)** Representative images illustrating the diameter comparison between the same vascular segments before and after CUS. **(d)** Statistics for vascular diameters before and after CUS for the same vascular segments. **(e)** Comparison of vascular diameter changes over the CUS period between stressed and unstressed mice. **(f)** Like e, but for the percentage of vascular diameter changes. d-f: control: n = 6/74; CUS: n = 6/74, mice/vascular segments. **(g)** Illustrating a rapid continuous line scan in a single vascular segment for the measurement of blood flow velocity. **(h)** Longitudinal comparison of blood flow velocities before and after CUS for the same vascular segments. **(i)** Changes in blood flow velocity over the period of CUS in stressed and unstressed mice. **(j)** Like (i), but for the percentage changes of blood flow velocity. h-j: control: n = 6/27; CUS: n = 6/27, mice/vascular segments).

We then sought to determine how individual vascular segments were impacted by chronic stress. To this end, we estimated vascular diameters and blood flow velocity from the same vascular segments before and after CUS. For mice under normal conditions, vascular diameter remained stable over time (Fig.3c, d). However, in the CUS mice, vascular segments presented decreased diameters (Fig.3c-d). As a result, after ten days of CUS, the vascular diameter in the CUS mice was reduced significantly more than the normal conditioned mice (Fig.3e, f). Accordingly, the overall vascular diameter in CUS mice was smaller than the normal mice (sFig3e). Like the vascular diameter changes, blood flow velocity was slower in most of the surveyed vascular segments after CUS induction, on average, reaching more than 300 µm per second slower than before CUS (Fig.3g-j). The reduction in blood flow velocity, along with the decreased microglial motility after CUS, is in line with the weakened motility of microglia processes as observed previously in mice with transient ischemia, wherein the decrease of blood flow correlated with the reduction of microglial motility ^30^. These data clearly indicate that CUS induces structural and functional dysregulations in the brain vasculature.

### 3.4 Microglial-vascular contact situations affect vascular alterations in depression

The parallel changes in microglial-vascular contact and vascular dysregulation following CUS prompted us to ask whether microglial-vascular contact dynamics influence vascular properties. To investigate this, using the data that imaged microglia and vasculature over one hour after CUS as described above (Fig.2a, b), we compared diameter changes in individual vascular segments that experienced various microglial contact dynamics over the same period. We found that in normally conditioned mice, no matter how vascular segments contact with microglia, they exhibited no significant diameter changes within one hour (Fig.4a, b). For the CUS mice, while the microglia uncontacted vessels exhibited no diameter changes, the microglia contacted vessels, especially those maintained contact within the observing one hour, showed a remarkable reduction in their diameters (Fig.4c, d). Additionally, vascular segments that lost contact with microglia also showed pronounced constriction (Fig.4c, d). The different effects of microglia on vascular diameter between mice with and without CUS resulted in distinct vascular diameter changes under the same microglia-vascular contact situations (Fig.4e-h). Collectively, these data indicate that in the stressed mice, vascular diameter was differentially modulated by microglia based on their contact situations with microglia. Here, vessels that are in contact with microglia are more likely to experience constriction than those that do not receive microglial contact.

**Figure 4.**
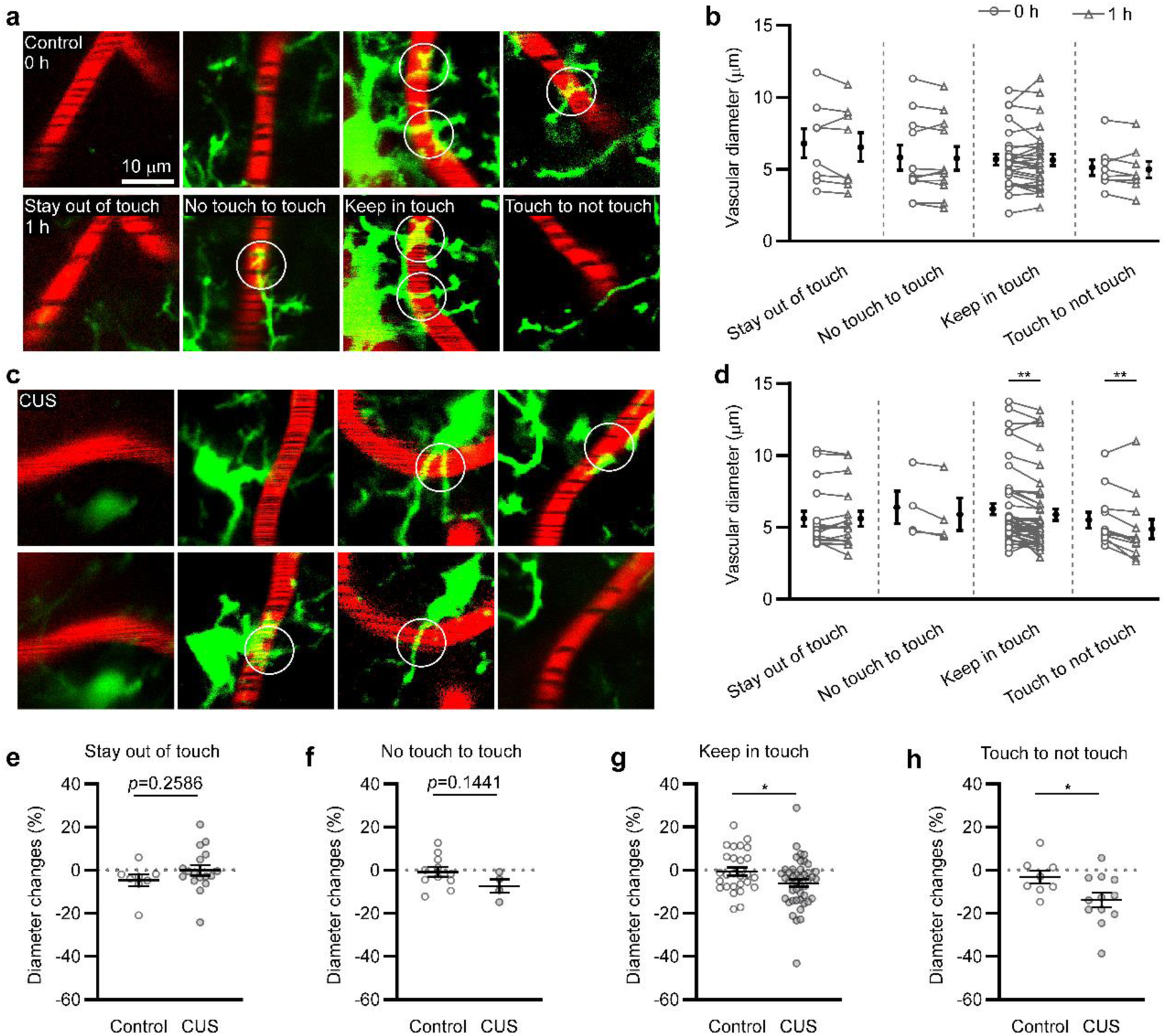
Microglial-vascular contact dynamics affect vascular diameter changes after CUS. **(a)** Representative images showing microglia vascular contact situations at two imaged timepoints (0 h and 1 h) in normal conditioned mice. White circles indicate contacts between microglia and vasculature. **(b)** Vascular diameter changes over 1 hour in unstressed mice. Based on microglia contact situations at the two timepoints, vascular segments were grouped as stay out of touch (n = 8 vascular segments), no touch to touch (n = 11), keep in touch (n = 27), touch to not touch (n = 8). **(c, d)** Like a and b, but for the analyzation of stressed mice. For vascular segments in stressed mice, stay out of touch (n = 17), no touch to touch (n = 4), keep in touch (n = 45), touch to not touch (n = 12). **(e-h)** Comparison of the vascular diameter changes between stressed mice and normally conditioned mice. The data source was the same as (b) and (d).

### 3.5 AT1R signaling in microglia mediates vascular changes under CUS

Given that microglia contact with vessels differentially affects vessels than those without contact with vessels, there might be different properties between microglia with and without vascular contact. To verify this, we first compared the morphology between microglia that contacted and uncontacted with the vasculature. In consistent with previous studies, when microglial somata were located on vessels, these somata were significantly bigger (sFig4, Table S1). However, the other morphological features showed no differences between microglia regardless of their contact situations with vessels (Table S1), suggesting mechanisms beyond microglial morphology underlie the differential regulation of vessels with and without microglial contacts.

As AngII and its type1 receptor – AT1R are involved in the renin-angiotensin system and act as vasoconstrictors ^36,37^, we next examined the expression of AngII and AT1R in microglia by combining RNAscope and microglia immunostaining (Fig.5d). We found that the overall expression of AngII and AT1R showed no significant differences between CUS mice and normal mice (Fig.5e, f). Interestingly, the spatial co-expression of AngII and AT1R was remarkably higher in CUS mice (Fig.5g). When analysg the expression of AngII and AT1R in microglia, we found that the expression of both AngII and AT1R was upregulated in microglia after CUS though their overall expression presented no differences compared with that in normal mice (Fig.5h, i). Accordingly, the spatial co-expression of AngII and AT1R after CUS was extremely present in microglia (Fig.5j). The CUS-induced upregulation of AngII and AT1R in microglia could enable microglia to mediate vasoconstriction. As vasoconstriction within a short time after CUS depends on the microglial-vascular contacts, the microglial expression of vasoconstrictors might also be different between microglia that are in contact and uncontract with vasculature. As expected, the overexpression of AT1R was only observed in vascular contacted microglia (Fig.5k, l), differentiating the vascular contacted microglia to induce local vasoconstriction.

**Figure 5.**
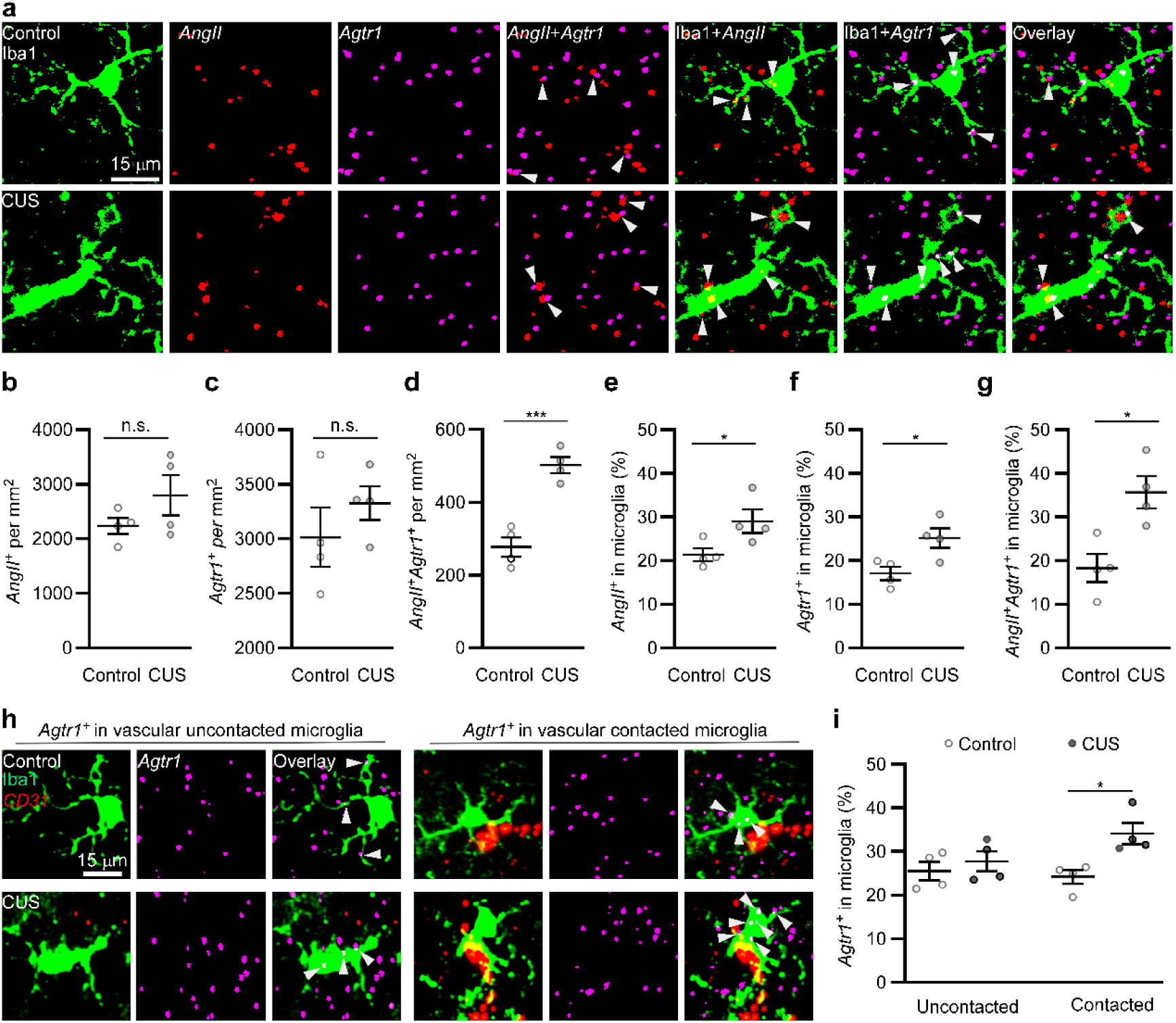
The mRNA expression of AngII and AT1R is upregulated in microglia. **(a)** Representative images showing immunofluorescence stained Iba1-positive cells, RNAscope labelled AngII and Agtr1. Arrows indicate the coexpression of AngII and Agtr1, the expression of AngII in Iba1-positive cells, the expression of Agtr1 in Iba1-positive cells and the co-expression of AngII and Agtr1 in Iba1-positive cells. **(b-g)** Statistical analysis for the expression of AngII and Agtr1, the co-expression of AngII and Agtr1, the percentile of expressed AngII in microglia, the percentile of expressed Agtr1 in microglia, and the percentage of conjugated AngII-Agtr1 in microglia. Control: n = 4 mice; CUS: n = 4 mice. **(h)** Representative images showing immunofluorescence stained Iba1 and RNAscope labelled mRNA expression of Agtr1 and CD31, with white arrows indicating expression of Agtr1 in Iba1-stained microglia. **(i)** Expression of Agtr1 in microglia with and without vascular contact (Control: n = 4 mice; CUS: n = 4 mice).

We next asked whether AT1R signaling in the CUS mice also contributed to other vascular pathologies and depression phenotypes. To answer this question, we pharmacologically blocked AT1R signaling in the chronically stressed mice via candesartan. We found that candesartan application prevented vasoconstriction in the CUS mice (Fig.6a, b). Meanwhile, normal vascular density and blood flow velocity were also preserved after CUS in the candesartan administrated mice (Fig.6c-g, sFig5a). In association with the beneficial effects of candesartan on vasculature, candesartan administration suppressed the microglia activation, showing comparable cell density and morphological properties to those in normal conditioned mice (Fig.6h-k, sFig.5b, c), suggesting crucial roles of AT1R in microglia activation under chronic stress. In parallel, microglial-vascular contacts remained normal in candesartan-applied CUS mice (Fig.6l-n). In line with these protective effects of candesartan on vessels and microglia under chronic stress, the candesartan administrated mice after CUS performed like normal conditioned mice, showing no obvious depression-like behaviours (Fig.6o-r), indicating critical roles of AT1R signaling in the development of depressive under chronic stress.

**Figure 6.**
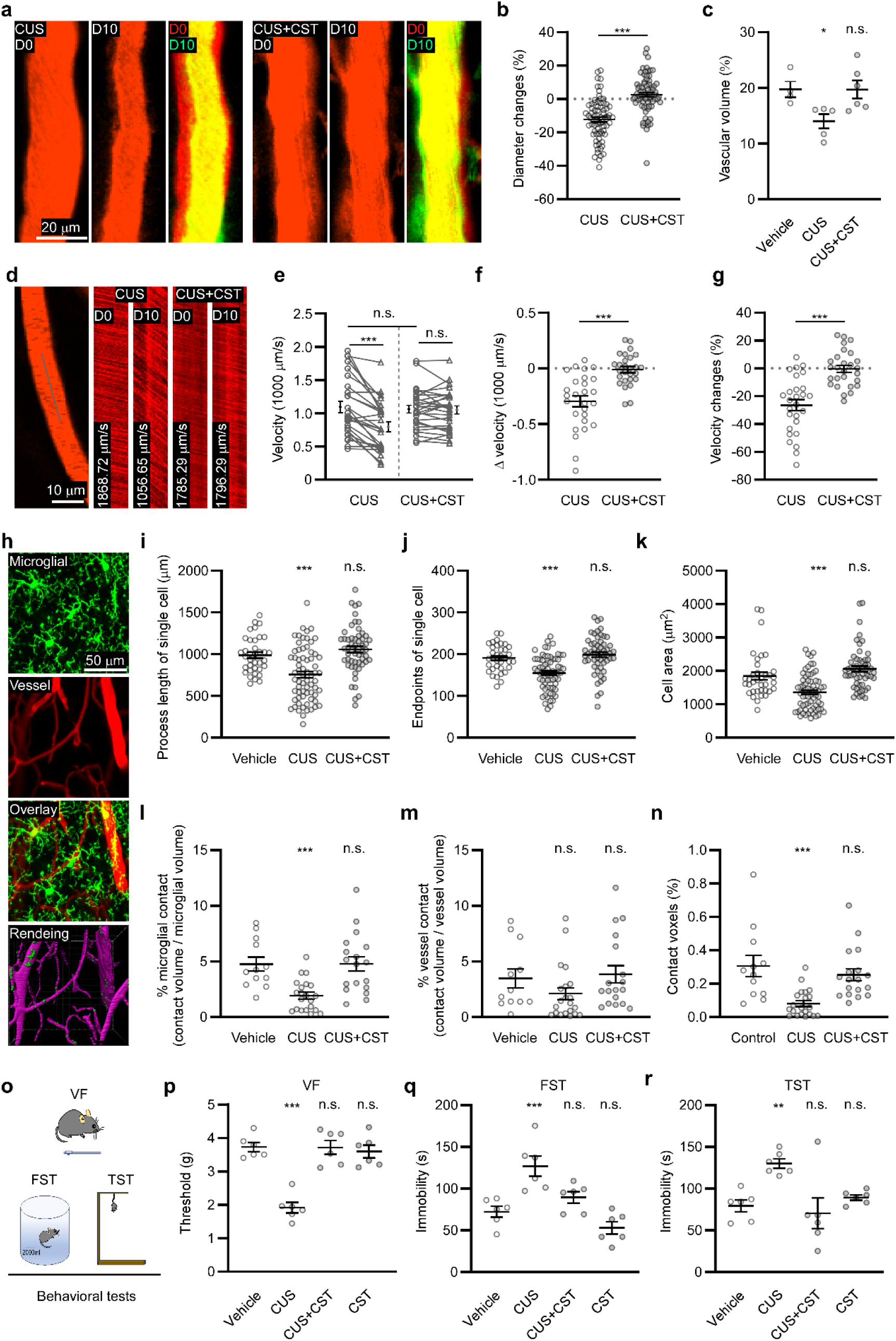
Effects of candesartan on vasculature, microglia, and depression-like behaviors. **(a)** Representative images showing the same vascular segments scanned before (D0) and after (D10) CUS. **(b)** Comparison of the vascular diameter changes between stressed mice with and without candesartan administration (CUS: n = 6/74; CUS+CST: n = 6/73, mice/vascular segments). **(c)** Quantification of vascular density (Vehicle: n = 4 mice; CUS: n = 5 mice; CUS+CST: n = 6 mice). **(d)** Exemplary measurement of blood flow velocity from the same vascular segment before and after CUS. **(e)** Statistics for blood flow velocity before and after CUS for the same vascular segments in mice with and without candesartan application. **(f)** Blood flow velocity changes induced by CUS. **(g)** Like (f), but for the percentile changes of blood flow velocity. e-g: CUS: n = 6/27; CUS+CST: n = 6/27, mice/vascular segments. **(h)** Representation of simultaneous imaging of microglia and blood vessels, and reconstructions of microglia (green) in contact with vasculature (magenta) using IMARIS. **(i-k)** Quantification of morphological parameters in single microglia. Vehicle: n = 4/36; CUS: n = 6/67; CUS+CST: n = 6/55, mice/cells. **(l)** Percentage of microglia area contacting with blood vessels in the total microglia areas. **(m)** The frequency of blood vessels in contact with microglia. **(n)** The percentage of colocalized microglial and vascular areas in the imaging region. l-n: Vehicle: n = 4/12; CUS: n = 6/21; CUS+CST: n = 6/18, mice/slice. **(o)** Illustrating behavioral tests. **(p)** Statistics for the paw withdrawal threshold of mice in the von Frey test. **(q)** Immobility time of mice in FST. **(r)** Immobility time of mice in TST. p-r: vehicle: n = 6 mice; CUS: n = 6 mice; CUS+CST: n = 6 mice; CST: n = 6 mice.

Together, these data show that chronic stress selectively enhances AT1R signaling in vascular contacted microglia, which enables the vascular contacted microglia to rapidly modulate vascular structure. The AT1R signaling also acts as a critical mediator for the general depression phenotypes, including microglia activation, pathological vascular alterations and depression-like behaviors.

### 3.6 Microglia activation under CUS advances vascular pathology and depression

The inhibition of AT1R activity associated with the suppression of microglia activation inspired us to probe whether inhibiting microglia activation under CUS could, in turn, suppress the AT1R activity and thus ameliorate vascular changes induced by CUS. For this, we pharmacologically suppressed microglia activation via minocycline. For the analyzation of microglia morphology and vascular diameter, we labelled vessels with lipid dye through intracardial perfusion as mentioned above, and acquired 3D images of microglia and vessels in the fixed brain slices. As expected, in the CUS mice, minocycline administration blocked microglia activation, displaying normal cell number and morphological properties (Fig.7a-d). The suppression of microglia activation was associated with no detectable CUS-induced alterations in the microglial-vascular contacts and vascular diameters (Fig.7f-h), indicating that normal microglial-vascular interactions were maintained under CUS. Furthermore, the stressed mice exhibited no depression-like behaviors with minocycline application (Fig.7i-l). These data suggest that microglia activation underlies the aberrant vascular-microglia interaction and abnormal vascular features under chronic stress.

**Figure 7.**
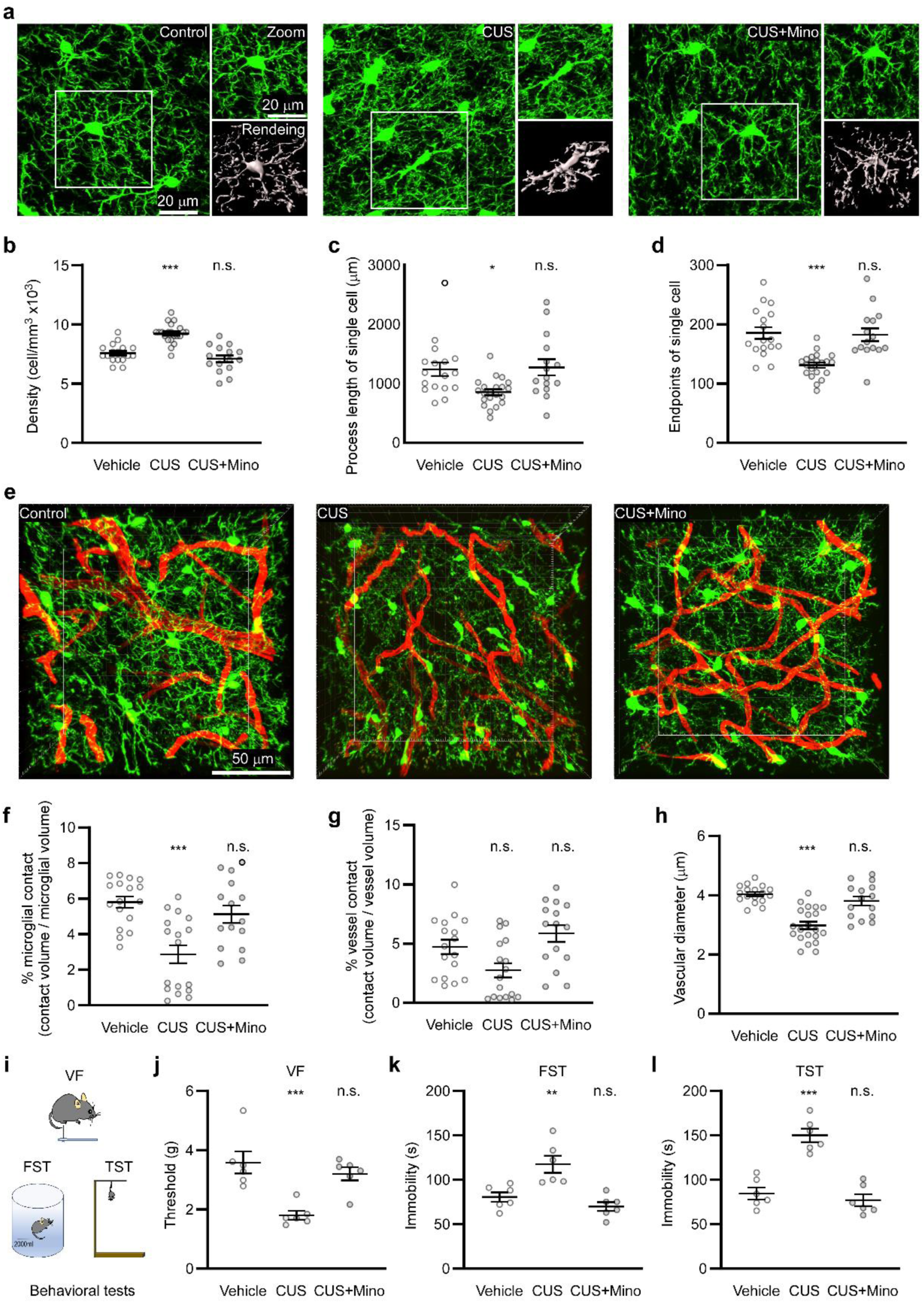
Inhibiting microglial activation by minocycline prevents CUS-induced vascular changes and depression-like behaviors. **(a)** Representative images showing microglia morphology and crops of single microglia (zoomed images) for the quantitative analyzation of morphological parameters in single microglia. **(b-d)** Statistical analysis for microglia density, microglial process length and the number of microglial endpoints. Control: n = 6/17; CUS: n = 7/21; CUS+Mino: n = 5/15, mice/slices. **(e)** Representative images showing microglia (green) and blood vessels (red) acquired from fixed brain slices. **(f-g)** Percentile comparison of microglia vascular contact area relative to microglial area (f) and vascular area (g) between mice with and without minocycline treatment. **(h)** The comparison of the averaged vascular diameter between mice with and without minocycline administration. f-g: control: n = 6/17; CUS: n = 7/21; CUS+Mino: n = 5/15, mice/slices. **(i-l)** Behavioral analyzation of mice in VF, FST and TST, and the perspective statistical results. vehicle: n = 6 mice, CUS: n = 6 mice; CUS+Mino: n = 6 mice.

Guided by the curious regulation of the chemokine fractalkine receptor – CX3CR1, on microglia activation ^38,39^, we next probed whether CX3CR1 deficiency was associated with a different outcome following chronic stress. For this, we compared microglial and vascular features between CX3CR1^GFP/+^ mice and the CX3CR1 deficient – CX3CR1^GFP/GFP^ mice. Before CUS, while microglia number and process length of single microglia showed no differences between mice with and without CX3CR1 deficiency, the terminal branches of microglia in CX3CR1 deficient mice were significantly less than CX3CR1^GFP/+^ mice (Fig.8a-d), indicating the lack of CX3CR1 led to altered microglia morphology, consistent with previously reported ^40^. Interestingly, after CUS, CX3CR1 deficiency alleviated the CUS-induced microglia activation, vasoconstriction and abnormal behaviour performances (Fig.8b-k). These data together suggest that CX3CR1 signaling is necessary for microglia activation under chronic stress, which further facilitates the vascular abnormality and depression-like behaviors.

**Figure 8.**
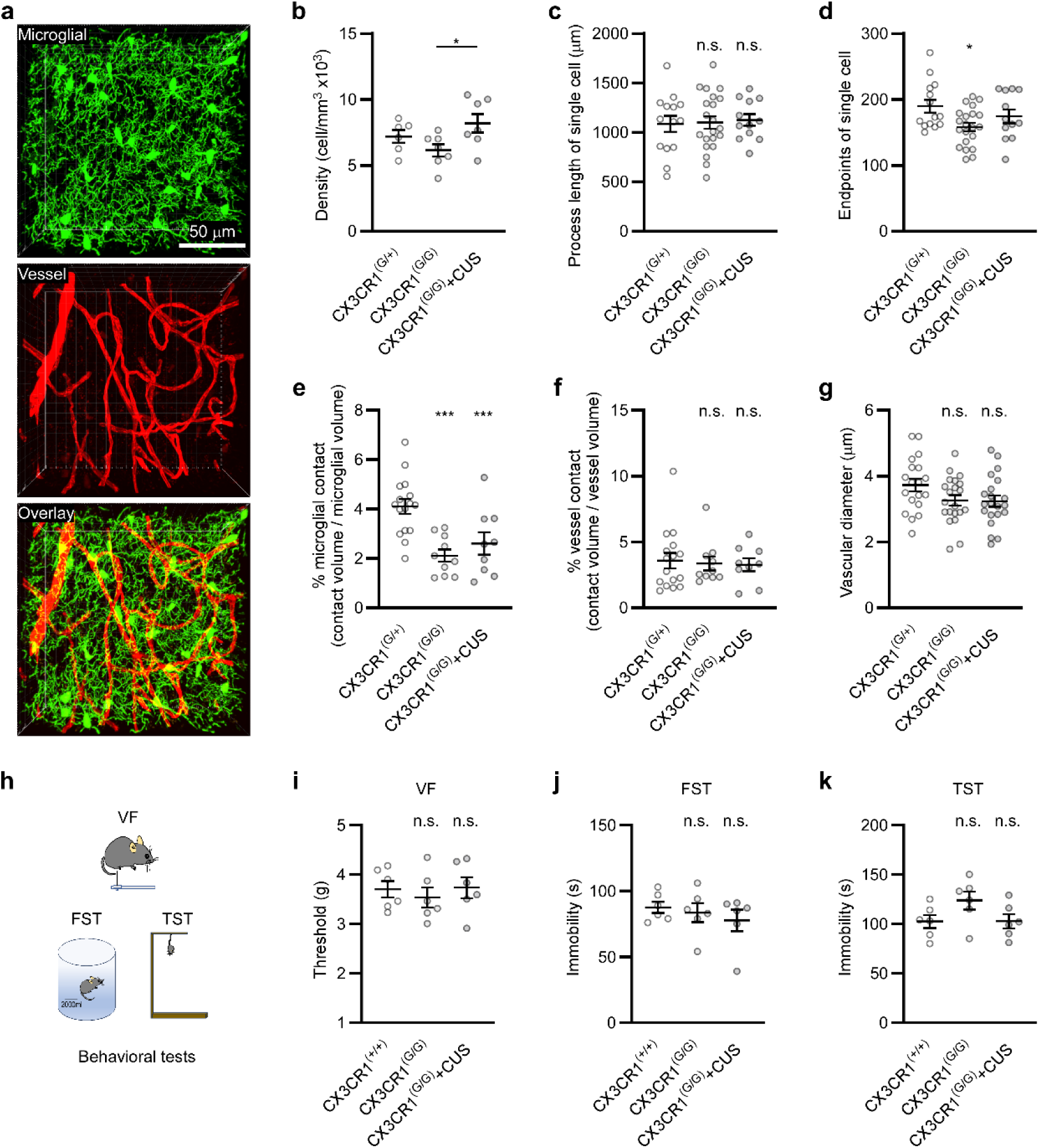
Knockout chemokine receptor CX3CR1 prevents depressive disorders under CUS. **(a)** Representative images showing microglia and blood vessels acquired from CX3CR1^GFP/GFP^ transgenic mice. **(b)** Comparison of microglia density between mice with and without CX3CR1 knockout after CUS. CX3CR1^G/G^: CX3CR1^GFP/GFP^; CX3CR1 ^G/+^: CX3CR1^GFP/+^. CX3CR1^G/+^ control: n = 6 mice; CX3CR1^G/G^ control: n = 7 mice; CX3CR1^G/G^ CUS: n = 7 mice. **(c-d)** Total length of microglial processes and the number of process endpoints in single microglia. CX3CR1^G/+^ control: n = 5/14; CX3CR1^G/G^ control: n = 7/21; CX3CR1^G/G^ CUS: n = 5/12, mice/cells. **(e-f)** Percentile comparison of microglia vascular contact area relative to microglia area and vascular area between mice with and without CX3CR1 knockout. CX3CR1^G/+^ control: n = 7/16; CX3CR1^G/G^ control: n = 5/10; CX3CR1^G/G^ CUS: n = 5/9, mice/slices. **(g)** Averaged vascular diameter in CX3CR1^G/+^ and CX3CR1^G/G^ mice. CX3CR1^G/+^ control: n = 7/19; CX3CR1^G/G^ control: n = 7/21; CX3CR1^G/G^ CUS: n = 7/21, mice/slices. **(h-k)** The VF, FST and TST tests in CX3CR1^+/+^ and CX3CR1^G/G^ mice with and without CUS experiences and the quantitative results. CX3CR1^+/+^ control: n = 6 mice; CX3CR1^G/G^ control: n = 6 mice; CX3CR1^G/G^ CUS: n = 6 mice.

Together, these results show that suppressing microglia activation under chronic stress via minocycline administration or CX3CR1 knock out prevents the pathological vascular changes and depression-like behaviors, supporting that microglia activation facilitates the development of depression under chronic stress.

## 4 Discussion

In this study, we have characterised the microglial-vascular interaction in depression induced by CUS and provided evidence for the involvement of their dysregulation in the pathogenesis of depression. We show that in CUS-induced depression, microglia exhibit reduced process motility and decreased contact area with vasculature. Meanwhile, the vasculature presents structural and functional pathologies. We find that the overall vascular compromises following CUS is accompanied by rapid vasoconstriction induced by vascular contacted microglia which present upregulated AT1R expression. We identify that AT1R singling together with microglia activation are key contributors to the pathological vascular alterations following CUS and other depression symptoms. Collectively, these findings indicate that CUS impairs the normal microglial-vascular interaction, which results from the enhanced microglial AT1R signalling and microglia activation, promoting vascular abnormalities and depression-like behaviors. This work offers novel insights into how aberrant microglial-vascular interactions contribute to depression under chronic stress.

Microglia are known to play crucial roles in depressive disorders ^14–16^. Much attention has been devoted to their interactions with neurons, which can lead to impaired functions in the neural network ^16,41–44^. However, the relationship between microglia and the vasculature has been largely overlooked to date. Although microglia-mediated inflammatory responses in depression may affect vascular structure and function, recent studies have reported that microglia directly contact with blood vessels and regulate cerebral blood flow ^19,20^. However, how microglia regulate vasculature under pathological contexts and how the microglial-vascular contact situations functionally relate to vascular modification remain poorly understood. In the present study, we show that chronic stress leads to a reduction in physical contact area between microglia and blood vessels, concomitant with decreased vascular density, vascular diameter and blood flow velocity resulted from the enhanced AT1R signaling in microglia. These effects of microglia on vascular structure and function appears contrary to observations in cerebrovascular injuries – where microglia migrate toward blood vessels and alleviate vascular impairments ^45–48^. These different changes in microglia-vascular contacts suggest that microglia exert distinct modulation on blood vessels in different pathological contexts.

Microglial diversity in gene expression, morphology and function has been increasingly reported. This heterogeneity has traditionally been attributed to differences in developmental stages, microglia activation status and brain regions ^49–52^. Recent evidence has shown the heterogeneous properties of microglia in the same brain region under the same physiological context ^19,53,54^. For example, in cortex, microglia in contact with vasculature exhibit bigger somata compared to those not in contact with vasculature ^19^. Our data show that within one hour in the stressed mice, vessels in contact with microglia display decreased diameter, which does not occur in vessels without microglia contact, indicating that microglia those contacted and uncontacted with blood vessels exhibit different regulation on vasculature. This finding highlights the functional divergence between different microglia subgroups in the same context.

The prevailing hypothesis regarding microglial regulation of vasculature in diseases emphasizes microglia activation induced inflammatory responses, which have also been proposed as the main mechanism underlying the reduction of blood perfusion in depression ^3,55–57^. If inflammation alone counted for the vascular alterations in depression, all vascular segments within the same pathological milieu would be expected to exhibit uniform changes. However, although nearly all observed vascular segments present decreased diameter following CUS, short-term post-CUS examinations reveal that vasoconstriction occurs preferentially at sites where vessels are in contact with microglia. This contact-dependent microglial regulation of vascular properties suggests the involvement of mechanisms beyond neuroinflammation.

Regarding the regulation of blood vessels by vascular contacted microglia, recent studies have shown that these cells can regulate vascular properties through P2RY12 signaling ^19,20,29^. Although AT1R is known as a vasoconstrictor that regulate blood pressure, the regulating mechanism is attributed to microglia activation associated inflammatory processes ^58–61^. In this study, our data suggest that AT1R signaling in microglia contributes to vascular alterations under chronic stress involving mechanisms beyond inflammation. Following CUS, we observed upregulated AT1R expression exclusively in vascular contacted microglia, enabling these microglia to mediate vasoconstriction at the microglial-vascular contact regions, as we observed the vascular diameter changes within one hour after CUS.

In addition to regulate the vascular diameter, AT1R signaling has been suggested to be involved in the pathogenesis of depression. Previous studies have demonstrated that chronic perfusion of AngII induces depression, suggesting that AngII-AT1R signaling contributes to the development of depression ^60,62,63^. However, whether depression is associated with a systematic increase in AngII levels remains unclear. Our findings indicate that in chronic stress-induced depression, AT1R activation and associated vasoconstriction occur without an overall elevation of AngII. Instead, AngII expression is selectively upregulated in microglia and presents more co-expression with AT1R, implying that vasoconstriction associated with depression may not be attributable to systemic AngII increases, but rather to microglia-specific AngII-AT1R signaling. Furthermore, our pharmacological interventions demonstrate that inhibiting AT1R activity restores normal vascular function and ameliorates depression-related alterations under chronic stress. These results highlight the critical roles of AT1R activation in mediating stress-induced depressive abnormalities and identify it as a key mechanistic factor in the pathogenesis of depression under chronic stress conditions.

The critical roles of both AT1R signaling and microglia activation in the induction of depression raise the question of what is the relationship between AT1R signaling and microglia activation. Our data from candesartan treatment indicate that AT1R signaling acts as an inducer of microglial activation, as previous studies suggested ^58–60^. Furthermore, our data show that inhibiting microglial activation prevents vascular alterations and depression-like behaviors following CUS, suggesting that microglia activation is necessary for vascular pathogenesis and the associated upregulation of AT1R signaling. Thus, AT1R signaling and microglia activation are likely to reinforce each other. Notably, the chemokine receptor CX3CR1 is required for microglial activation and stress-induced responses following CUS as demonstrated in the current study and other studies ^38,39^. Therefore, CX3CR1 signaling may serve as a critical intermediary between AT1R signaling and microglia activation, whereby upregulated AT1R signaling drives microglial activation, and in turn, activated microglia promotes the upregulation of AT1R signaling, thereby exacerbating vascular dysregulation and ultimately leading to depression. However, further investigation is required to elucidate whether and how the detailed mechanistic interplay between AT1R signalling and microglial activation differs between vascular-contacted microglia and vascular-uncontacted microglia. Additionally, future studies should examine how the other vasoconstriction related genes, particularly those expressed in microglia, contribute to the abnormalities of microglial-vascular interaction and the development of depression under CUS.

## 5 Conclusion

In summary, we have characterized the microglial-vascular interaction alterations and the associated vascular pathologies in a mouse model of depression induced by CUS, identified the selective upregulation of AT1R signaling in microglia, and elucidated that both AT1R signaling and microglia activation are critical underlying mechanisms regulating microglial-vascular interactions and the depression-like behaviors. Together, our findings provide evidence that abnormal microglial-vascular interactions induced by elevated microglial AT1R signaling and microglia activation promote the development of depression under chronic stress.

## Ethics statement

All animal experiments were performed in strict accordance with the guidelines established by the European Community (2010/63/EU) and the ARRIVE guidelines, and was approved by the Institutional Ethics Committee of the School of Basic Medical Sciences, Lanzhou University (Licence reference number: lzujcyxy20231203, 02 December 2023).

## CRediT authorship contribution statement

**Hang Gao:** Methodology, Investigation, Formal analysis, Data curation, software, Visualization, Writing – review & editing. **Zeyuan Ding:** Methodology, Investigation, Data curation, Writing – review & editing. **Biao Xu:** Methodology, Resources, Writing – review & editing. **Yuqing Wu:** Validation, Data curation, Writing – review & editing. **Meizhen Zhu:** Data curation, Writing – review & editing. **XiaoYue Zhang:** Formal analysis, Writing – review & editing. **Fujian Qi:** Methodology, Resources, Writing – review & editing. **Junru Liu:** Methodology, Resources, Writing – review & editing. **Quan Fang:** Methodology, Resources, Supervision, Writing – review & editing. **Yanli Ran:** Writing – original draft, Formal analysis, Visualization, Resources, Project administration, Supervision, Conceptualization, Funding acquisition.

## Acknowledgments and funding sources

We thank Hao Gao and Yongrong Qiu for their technical helps. We thank New Medical Science Innovation Platform of Lanzhou University for providing instrument supports. This work was funded by the National Natural Science Foundation of China (No.32200810) and the “Double First-Class” Research Start-up Funds of Lanzhou University (561120203).

## Data availability

All data presented in this study will be available upon the publication of this manuscript.

